# Neighborhood regulation by lncRNA promoters, transcription, and splicing

**DOI:** 10.1101/050948

**Authors:** Jesse M. Engreitz, Jenna E. Haines, Glen Munson, Jenny Chen, Elizabeth M. Perez, Michael Kane, Patrick E. McDonel, Mitchell Guttman, Eric S. Lander

## Abstract

Mammalian genomes are pervasively transcribed to produce thousands of spliced long noncoding RNAs (lncRNAs), whose functions remain poorly understood. Because recent evidence has implicated several specific lncRNA loci in the local regulation of gene expression, we sought to determine whether such local regulation is a property of many lncRNA loci. We used genetic manipulations to dissect 12 genomic loci that produce lncRNAs and found that 5 of these loci influence the expression of a neighboring gene in *cis*. Surprisingly, however, none of these effects required the specific lncRNA transcripts themselves and instead involved general processes associated with their production, including enhancer-like activity of gene promoters, the process of transcription, and the splicing of the transcript. Interestingly, such effects are not limited to lncRNA loci: we found similar effects on local gene expression at 4 of 6 protein-coding loci. These results demonstrate that ‘crosstalk’ among neighboring genes is a prevalent phenomenon that can involve multiple mechanisms and *cis* regulatory signals, including a novel role for RNA splicing. These mechanisms may explain the function and evolution of some genomic loci that produce lncRNAs.

---

Mammalian genomes are pervasively transcribed^1–4^ to produce thousands of spliced and polyadenylated long noncoding RNAs (lncRNAs)^5–7^, most of whose functions remain unknown. Recent evidence shows that some genomic loci that produce lncRNAs regulate the expression of nearby genes. In a few cases – including XIST^8,9^, HOTTIP^10^, and others^11–13^ – lncRNAs have been shown to recruit regulatory complexes to influence local gene expression. It has been suggested that many other lncRNAs similarly act as local regulators^14–16^. Such local regulatory functions might help explain the observation that lncRNA expression is often correlated with the expression of nearby genes^6,7,17–19^. However, such correlations could alternatively be due to other mechanisms: for example, gene promoters have been proposed to have dual functions as enhancers^20–25^, and the process of transcription *per se* has been proposed to contribute to gene regulation by recruiting activating factors or remodeling nucleosomes^26–30^. It has been challenging to identify and distinguish among local functions mediated by lncRNA promoters, the process of transcription at lncRNA loci, or the RNA transcripts themselves^31^.

We set out to identify lncRNA loci that participate in local gene regulation and to dissect the mechanisms that mediate these regulatory effects. To begin, we developed a genetic approach to distinguish between (i) primary effects on expression of nearby genes resulting from direct local functions of the lncRNA locus and (ii) secondary effects on nearby genes resulting from indirect downstream consequences of the lncRNA acting elsewhere in the cell (**Fig. 1a**, **Note S1**). We generated clonal cell lines carrying heterozygous genetic modifications at lncRNA loci and compared the expression of genes within 1 megabase (neighboring genes) on the *cis* and *trans* alleles (*i.e.*, on the modified and unmodified homologous chromosomes) in the same cells (**Fig. 1b**). Changes in neighboring gene expression that involve only the *cis* allele likely result from local functions, while changes that involve both the *cis* and *trans* alleles likely result as downstream effects of non-local functions (**Note S1**). We performed genetic modifications in 129/Castaneus F1 hybrid mouse embryonic stem cells (mESCs) that contain ~1 polymorphic site every 140 basepairs (bp), enabling us to distinguish the two alleles using RNA sequencing (**Fig. S1**), and we checked for consistency between knockouts on each genetic background to control for potential haplotype-specific effects (**Note S1**).

**Fig. 1.**
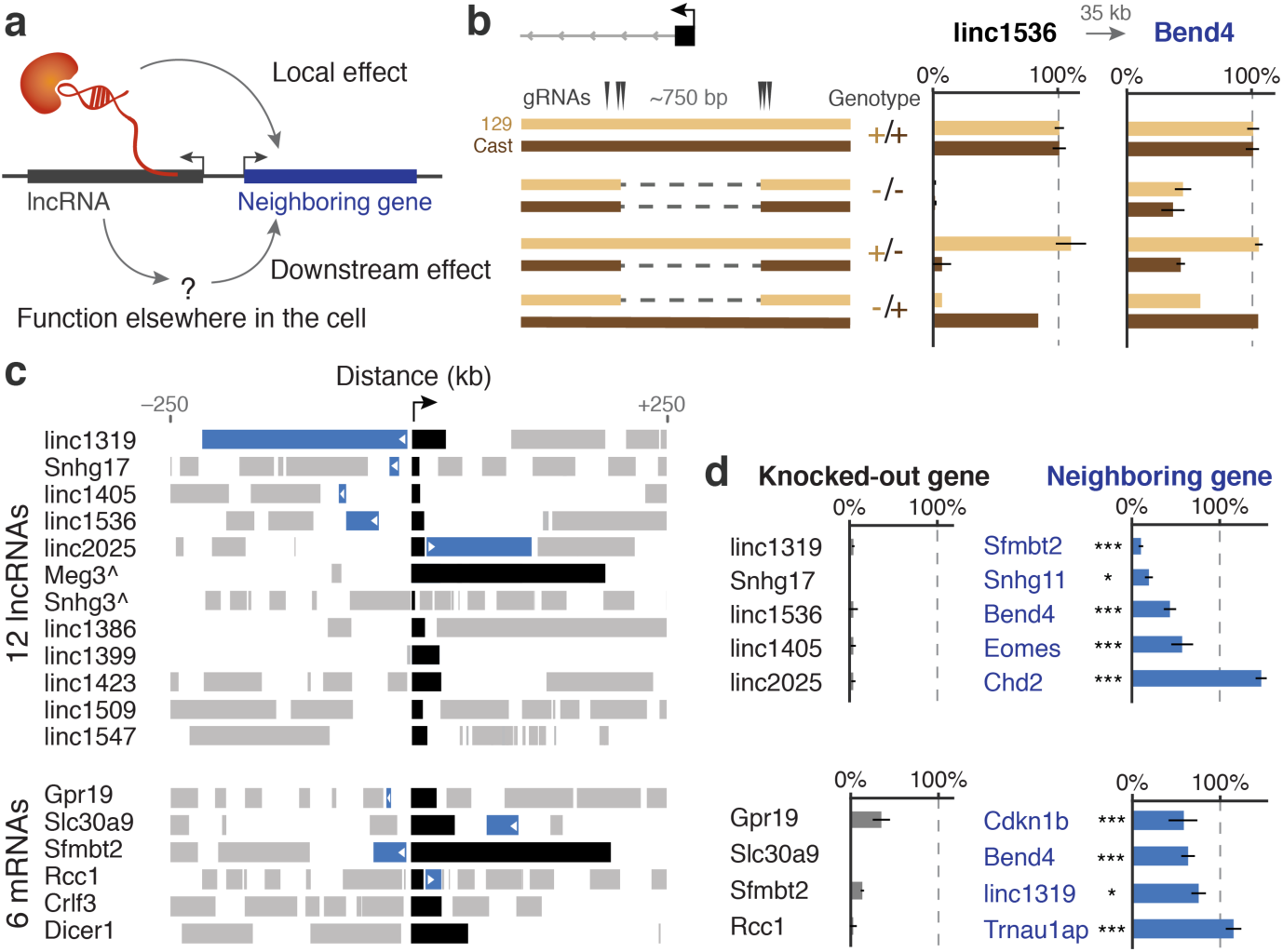
Many lncRNA and mRNA loci influence the expression of neighboring genes. (**a**) Knocking out a gene promoter (black) might affect the expression of a neighboring gene (blue) through local or non-local functions of the locus. (**b**) Knockout of the linc1536 lncRNA promoter. DNA from individual mESC colonies (left) was genotyped to identify 4 homozygous (−/−) and 5 heterozygous knockouts (+/− and −/+). Allele-specific RNA expression of linc1536 and a neighboring gene, Bend4, is normalized to the average of 81 control clones (+/+). Error bars: ± 95% confidence interval (CI) for the mean. **(c)** Gene neighborhoods for 12 lncRNAs and 6 mRNAs, oriented so each knocked-out gene is transcribed in the positive direction. Blue genes show allele-specific changes in RNA expression upon knocking out the promoter of the black gene; gray genes do not. ^The Meg3 and Snhg3 loci show effects attributable to read-through transcription (see Note S3). We examined all genes within 1 Mb of the lncRNA locus (Figs. S4 and S5); here we show genes within 250 kb, which includes all genes affected upon promoter knockout. **(d)** Promoter knockouts eliminate the expression of the targeted gene (black) and, in 9 loci, affect the expression of a neighboring gene (blue). Bars: Average RNA expression on knockout compared to wild-type alleles (see Methods). Error bars: 95% CI for the mean. *: FDR < 10%. ***: FDR < 0.1%. The FDR does not necessarily correlate with the size of the CI because each gene has a different variance in the wild-type controls (not pictured).

We dissected the *cis*-regulatory functions of 12 lncRNA loci whose RNA transcripts show preferential localization to the nucleus and span a range of abundance levels (for selection criteria, see Methods, **Fig. S2**, **Table S1**). For each locus, we co-transfected mESCs with Cas9 and multiple guide RNAs to delete ~600-1000 bp centered on the transcription start site (TSS) (**Fig. 1b**), reducing the transcript levels of the targeted allele by an average of 94% (**Table S1**). To identify loci that locally regulate gene expression, we looked for allele-specific changes in the expression of genes that neighbor the knocked-out locus (**Note S2**, see Methods). In total, 5 of the 12 lncRNA promoter knockouts significantly affected the expression of a neighboring gene at a false discovery rate (FDR) of < 10%, including both activating and repressive effects (**Fig. 1c**,**d**, **Note S3**, **Fig. S3**). For each locus, only a single neighboring gene showed a significant allele-specific change, and in each case the affected gene was located immediately adjacent to, and within 5-71 kb of, the knocked-out promoter (**Fig. 1c**, **Fig. S4**). Thus, a substantial fraction of these lncRNA loci influence the expression of an immediately neighboring gene.

We considered whether such effects were specific to lncRNA loci, or whether similar effects might be seen for mRNA loci. We therefore deleted the promoters of 6 protein-coding genes (**Fig. S2**, see Methods for selection criteria). Interestingly, knockouts for 4 of these protein-coding loci also altered the expression of a neighboring gene in an allele-specific manner (**Fig. 1c**,**d**, **Fig. S5**), including several reciprocal effects on the expression of neighboring lncRNAs. For example, knocking out the promoter of Sfmbt2, which itself was affected by promoter knockout at linc1319, in turn affected the expression of linc1319 (**Fig. 1d**).

These results suggest that both non-coding and coding loci participate in a broader network of direct regulatory connections to neighboring genes. Such connections might partly explain the observed correlations in gene expression within local neighborhoods^18,32–34^. The mechanisms underlying such *cis* effects in endogenous gene loci remain relatively unexplored and could in principle involve (i) DNA regulatory elements in gene promoters; (ii) the process of transcription; or (iii) the RNA transcripts themselves (**Fig. 2a**). Relevant to the first possibility, recent studies have highlighted that gene promoters can act as enhancers in plasmid-based reporter assays^20–22,24,40^, frequently contact one another in the nucleus^20^, and, in a handful of loci, appear to directly regulate a neighboring gene^23,25,35^.

**Fig. 2.**
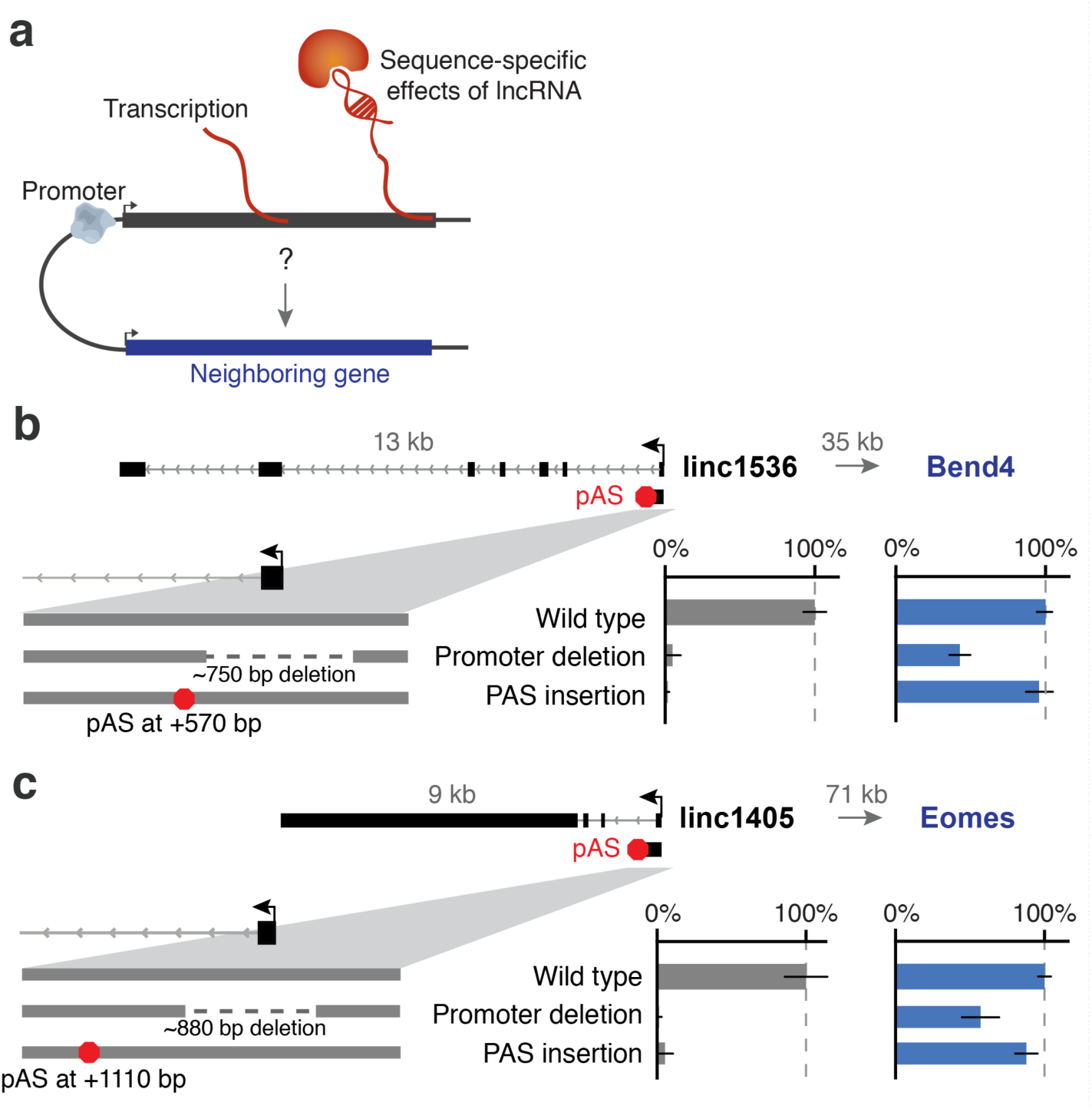
Some lncRNA promoters have enhancer-like activity. **(a)** Mechanisms by which a lncRNA locus (black) might regulate a neighboring gene (blue). **(b)** Average expression on knocked-out alleles compared to controls for linc1536 and the adjacent gene Bend4 upon deleting the linc1536 promoter or inserting a pAS. Linc1536 is encoded 35 kb away from Bend4. Linc1536 expression levels are assessed based on SNPs in the last three exons. Bars: Average RNA expression on 2+ knockout alleles compared to wild-type alleles (see Methods, Table S1). Error bars: 95% CI for the mean. Gray arrow indicates distance from the lncRNA promoter to the neighboring gene. (**C**) Allele-specific expression of linc1405 and an adjacent gene, Eomes, upon deleting the linc1405 promoter or inserting a triple pAS. Additional loci are presented in Fig. S8.

To begin to distinguish among these possible mechanisms, we inserted early polyadenylation signals (pAS), 0.5-3 kb downstream of each TSS, that eliminated the production of most of the RNA while leaving the promoter sequence intact (**Fig. 2**, **Fig. S6**, see Methods). We engineered pAS insertions that reduced RNA expression at 4 lncRNA loci and 2 mRNA loci (see **Note S4** for remaining loci), and examined their effects on the expression of neighboring genes.

At 5 of 6 loci where promoter deletion affected the expression of a neighboring gene, insertion of a pAS had no effect. For example, whereas deleting the promoter of the linc1536 locus reduced by 57% the expression of the adjacent Bend4 gene, insertion of a pAS into the first intron of linc1536 (~570 bp downstream of the TSS in this ~13kb locus) had no effect on Bend4 expression despite eliminating expression of the spliced linc1536 RNA (>99% reduction in levels of downstream exons) (**Fig. 2b**). This indicates that the full ~2 kb lncRNA is not required for Bend4 activation. In fact, the shortened ~570 bp transcript upstream of the pAS insertion is also unlikely to be required inasmuch as deep sequencing of the transcriptome revealed very little RNA derived from the sequence upstream of the pAS insertion (< 10% compared to the wild-type allele, **Fig. S7**) – perhaps because the pAS prevents RNA splicing, which may dramatically reduce transcriptional activity in the modified locus^36–39^. Therefore, the *cis* effect is likely mediated by DNA regulatory elements in the ~750 bp knocked-out promoter-proximal region.

Similar observations in 2 other lncRNA loci (linc1405, Snhg17) and 2 mRNA loci (Gpr19, Slc30a9) (**Fig. 2b**,**c**, **Fig. S8a-c**) suggest that the promoter-proximal sequences of many genes activate the expression of a neighbor. Although the promoters in these loci would not be classified as “enhancers” based on their chromatin state (*e.g.*, H3K4me3/H3K4me1 ratios^40^), they are marked by H3K27ac histone modifications (**Fig. S8d**), bound by mESC transcription factors (**Fig. S8d**), and are located in close spatial proximity to their neighboring target genes (**Fig. S8e,f**), suggesting that these gene promoters may affect expression of neighboring genes through mechanisms similar or identical to enhancers^20,41,42^. We note that the pAS insertion experiments do not rule out the possibility that these local regulatory effects might involve weak promoter-proximal transcription, similar to eRNA transcription at enhancers^42^.

While most of the local regulatory effects appeared to be mediated by promoter-proximal DNA elements, we identified one locus (linc1319) where promoter deletion and pAS insertion both substantially reduced the expression of a neighboring gene (Sfmbt2) (**Fig. 3a**). To dissect the regulatory mechanism, we first sought to determine whether the activation of Sfmbt2 is mediated by a sequence-specific function of the linc1319 RNA transcript or instead by the process of transcription *per se*. To test the first possibility, we knocked out each of the 3 downstream exons and 3 introns. We found that none of these deletions impaired Sfmbt2 activation (**Fig. 3a**). To the contrary, one of these deletions (removing 19.2 kb from the first intron) unexpectedly led to a ~5-fold *up-regulation* of linc1319 and a ~15-fold *increase* in Sfmbt2 expression (**Fig. 3a**–**b**, **Note S5**). These observations suggest that the activation of Sfmbt2 does not require unique sequences or structures in the linc1319 RNA transcript itself and instead may depend on the amount of transcription in the linc1319 locus. To test this possibility, we engineered pAS insertions at five different locations in the first exon or intron and found that increasing distance of the pAS from the linc1319 TSS (+40 bp to +15 kb, **Fig. 3a**) led to increased activation of Sfmbt2. These data indicate that the linc1319 locus activates Sfmbt2 through a mechanism that responds to the amount of transcription but does not require specific elements in the mature linc1319 RNA transcript. For example, the mechanism might involve the recruitment of transcription-associated factors such as pause release factors or chromatin regulators acting on the nearby Sfmbt2 promoter (**Fig. 3c**, **Note S6**).

**Fig. 3.**
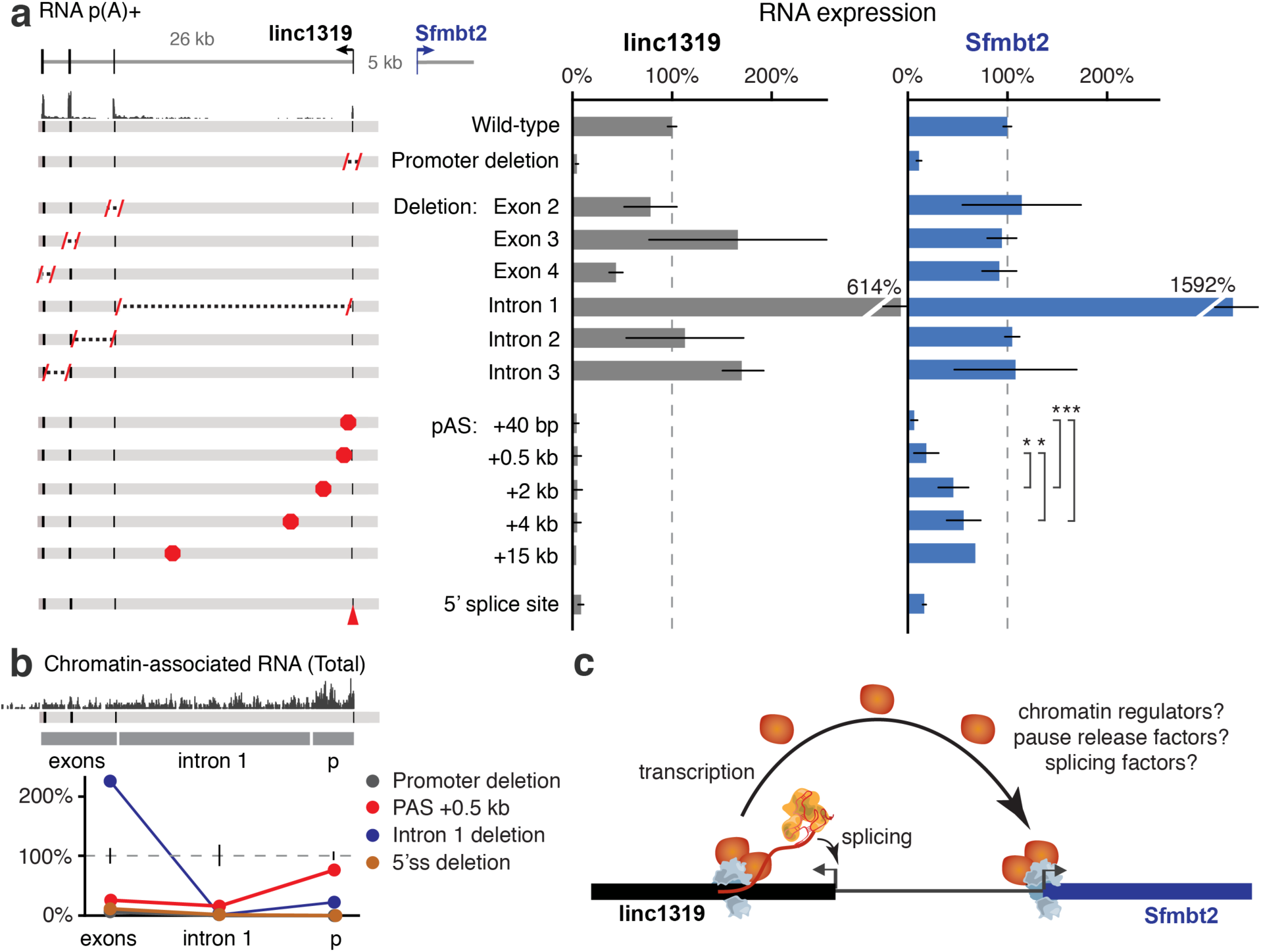
Transcription and splicing of linc1319 activates the expression of Sfmbt2. (**a**) (Left) Diagram of genetic manipulations in the linc1319 locus. (Right) Average RNA expression on knocked-out alleles compared to controls. Each bar presents data from 2+ independent clones, except for pAS at +15 kb, for which we obtained only 1 clone (see Table S1). Error bars: 95% CI for the mean. Error bars for Intron 1 deletions extend beyond the *y*-axis maximum. For linc1319, allele-specific RNA expression was measured using SNPs in exons 2-4; for the exon deletions, the deleted exon was excluded from this calculation. Sfmbt2 expression for pAS insertions was compared using a *T*-test: *p* < 0.05 (*) or < 0.01 (**). (**b**) Chromatin-associated total RNA – a proxy for transcriptional activity – for selected genetic manipulations. Raw read coverage at top depicts wild-type cells. The coverage of allele-informative reads in knockouts versus wild-type controls is averaged across 3 regions: the promoter-proximal 2.5 kb (“p”), the rest of intron 1, and the region downstream of intron 1 (“exons”). Error bars: 95% CI for the mean of 5 control clones. (**c**) Model for how transcription in the linc1319 locus activates the expression of Sfmbt2 by recruiting chromatin regulators, pause release factors, or splicing factors. Splicing of linc1319 further activates linc1319 transcription, leading to enhanced activation of Sfmbt2.

Because promoter-proximal splice sites and the process of splicing can enhance transcription-in some cases by as much as 100-fold^36–39^ - we tested whether the splicing of linc1319 affected transcription of linc1319 and Sfmbt2. Upon deleting the 5’ splice site of the first intron of linc1319 (**Fig. S9**), we observed a 93% reduction in the levels of chromatin-associated intronic RNA (a proxy for linc1319 transcription), a 92% reduction in the levels of the mature linc1319 transcript, and an 85% reduction in Sfmbt2 expression (**Fig. 3a**,**b**). These data demonstrate that the presence of the 5’ splice site of linc1319 is important for Sfmbt2 activation, revealing a novel role for splice signals in regulating neighboring genes. One possibility is that splicing promotes transcriptional activity in the linc1319 locus, which in turn recruits transcription-associated factors to the Sfmbt2 promoter; these activating factors may include the splicing complexes themselves (**Fig. 3c**). We note that in this model the linc1319 RNA transcript is in fact required for Sfmbt2 activation (splicing involves direct interactions between the spliceosome and the nascent transcript), yet the mechanism does not appear to depend on the precise sequence of the RNA beyond the presence of 5’ and 3’ splice signals.

In summary, genetic dissection of 12 lncRNA loci and 6 mRNA loci revealed that both protein-coding and noncoding loci often regulate gene expression in their local neighborhoods through general mechanisms associated with active transcription (**Fig. S10**). We did not identify any lncRNA loci in which local effects are mediated by sequence-specific functions of the lncRNA transcript. Instead, in most of the cases we studied, these effects are mediated by enhancer-like functions of H3K4me3-marked promoters, blurring the distinction between genomic elements classified as “enhancers” and “promoters” based on chromatin state^41,42^. As seen at one locus, the processes of transcription and splicing can also contribute to *cis* regulatory functions, perhaps by increasing the local concentration of transcription-associated factors. Notably, other co-transcriptional processes – including pause release, polyadenylation, and termination – involve distinct sets of regulatory factors that can feed back to modulate transcriptional activity^43–46^, suggesting that additional sequence signals associated with transcription might also contribute to *cis* effects in some loci. Together, these mechanisms enable local networks of regulatory connections, or ‘crosstalk,’ among active genes—including coding and non-coding loci. This crosstalk may explain in part the previously observed correlations in the expression of genes within local neighborhoods^7,18,32,33^. The properties of these *cis* regulatory connections – such as the mechanisms for specificity of regulation or the potential for cooperative dynamics of gene activation – represent key areas for future investigation.

While these mechanisms are present at both protein-coding and noncoding loci, they have particularly important implications for understanding the function and evolution of the genomic loci that produce lncRNAs. In loci where a promoter acts as an enhancer, RNA transcripts may arise as non-functional byproducts of an active *cis* regulatory element^25^. In loci where co-transcriptional processes have *cis* regulatory functions, the nascent transcripts themselves might contribute through mechanisms like splicing that require little RNA-sequence specificity. These possibilities are particularly intriguing in light of the observation that most lncRNA transcripts are not conserved across mammalian species^47–50^. Indeed, 65% of the 307 lncRNAs expressed in mESCs are “mouse-specific” (no syntenic transcript found in human, rat, or chimp pluripotent stem cells, **Fig. 4a**, see Methods)^50^, and the evolutionary conservation of these loci point to some candidates that may have *cis* regulatory functions independent of the lncRNA itself (**Fig. 4a**, **Fig. S11a**). As one exemplary category (see **Note S7** for others), 11 mouse-specific lncRNAs appear to have evolved from ancestral regulatory elements: the sequence serving as a promoter in mouse is adjacent to the same genes in mouse and human and corresponds to a conserved DNA element marked in human embryonic stem cells by a chromatin signature associated with enhancers (**Fig. 4b**–**d**, **Fig. S11b,c**, see Methods). These sequences may have conserved roles as *cis* regulatory elements, although not as lncRNA promoters. Because the lncRNA transcripts in these loci are not conserved, it is possible that some may represent non-functional byproducts of their promoters (**Fig. 4e**)^25^. Alternatively, some of these transcripts might contribute to *cis* functions without evolving specific RNA sequences or structures (**Fig. 4e**). These possible models are distinct (not necessarily mutually exclusive) from those in which lncRNAs function through sequence-specific RNA domains, and may explain the functions and evolution of an important subset of noncoding transcripts in mammalian genomes.

**Fig. 4.**
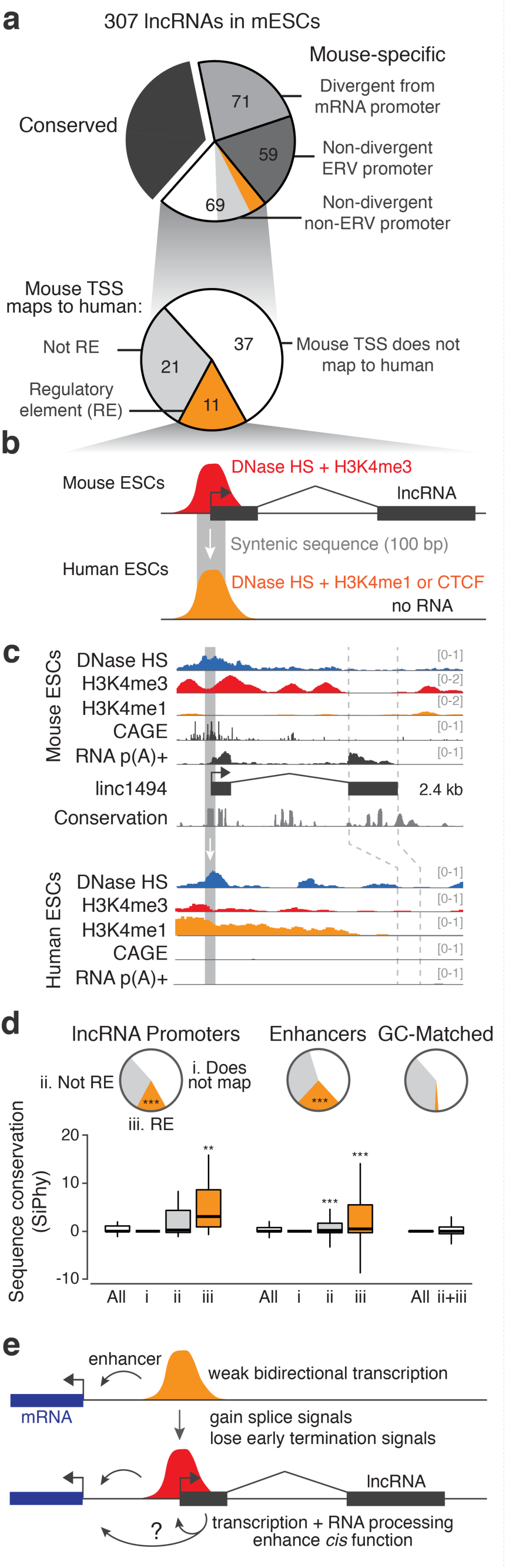
Evolutionary conservation of mESC lncRNAs and their promoters. **(a)** Classification of 307 lncRNAs expressed in mESCs. “Conserved” transcripts are those that show significant evidence of capped analysis of gene expression (CAGE) data and/or p(A)+ RNA in syntenic loci (see Methods). Divergent: initiating within 500 bp of an mRNA TSS, on the opposite strand. ERV: endogenous retroviral element (see Note S7). **(b)** Of 32 mouse-specific intergenic lncRNAs whose promoters map to syntenic sequence in human, 11 have promoters that correspond to putative DNA regulatory elements marked by DNase hypersensitivity (HS) in hESCs. (**c**) Example of a mouse-specific lncRNA (linc1494) whose promoter corresponds to a putative enhancer in hESCs. Conservation: PhastCons. (**d**) Sequence-level conservation of the promoters of mouse-specific lncRNAs, a random set of enhancers (matched to lncRNA promoters by H3K27ac signal in mESCs), and random intergenic regions (matched to lncRNA promoters by GC content). Positive SiPhy score indicates evolutionary constraint on functional sequences^51^. Categories and colors match pie chart in part (a). lncRNA promoters and enhancers in mouse are significantly enriched for corresponding to human regulatory elements (***: *P* < 10^−10^, Chi-squared test versus GC-matched). These sequences show elevated sequence-level conservation compared to GC-matched regions that map to human sequences (ii+iii) (**: *P* < 0.01, ***: *P* < 0.001, Mann-Whitney U-Test). Whiskers represent data within 1.5 × the interquartile range of the box. **(e)** Model for evolution of lncRNAs from pre-existing enhancers, which often initiate weak bidirectional transcription to produce eRNA^42^. Mutational patterns associated with transcription-coupled repair may favor the appearance of splice signals and loss of polyadenylation signals, promoting the neutral evolution of spliced transcripts^52^. In some cases, transcription, splicing, or other RNA processing mechanisms may feed back and contribute to the *cis* regulatory function of the promoter, producing a lncRNA as a byproduct.

While these models are attractive, it is important to note that the existence of these *cis* regulatory functions in a locus does not necessarily exclude other functions mediated by the RNA transcript itself. Indeed, we show that mRNA loci can both regulate neighboring genes in *cis* and produce mRNA transcripts, and the conserved Snhg17 locus serves as a host gene for snoRNAs in addition to activating a neighboring gene through the enhancer-like activity of its promoter (**Fig. S8a**). Similarly, other lncRNA transcripts in loci dissected here may also have functions in *trans* or in cellular contexts that we have not characterized. Nevertheless, it is clear that a full accounting of the functions of noncoding transcription will require further dissection not only of lncRNAs but also of the molecular mechanisms by which promoters, transcription, and RNA processing coordinate gene expression in local neighborhoods.

## Acknowledgements

We thank Shari Grossman, John Rinn, Moran Yassour, Phil Sharp, Laurie Boyer, Millie Ray, Charlie Fulco, Mathias Munschauer, Tim Wang, and Nir Friedman for discussions; Alex Shishkin for technical advice and reagents; and Jason Flannick for computational tools. J.M.E. is supported by the Fannie and John Hertz Foundation and the National Defense Science and Engineering Graduate Fellowship. This work was supported by funds from the Broad Institute (E.S.L.), the NIH Director’s Early Independence Award (DP5OD012190 to M.G.), the Edward Mallinckrodt Foundation (M.G.), the Sontag Foundation (M.G.), and the Searle Scholars Program (M.G.).

## Author contributions

J.M.E., M.G., and E.S.L. conceived and designed the study. J.M.E., J.E.H., G.M., and P.E.M. developed experimental protocols. J.M.E., J.E.H., G.M., E.M.P., and M.K. performed experiments. J.M.E. developed computational tools and performed data analysis. J.M.E. and J.C. performed evolutionary analysis. J.M.E. and E.S.L. wrote the manuscript with input from all authors.

